# Small but mitey: long-read assembly of a streamlined mite genome from contaminated host plant sequencing data

**DOI:** 10.1101/2024.05.17.594625

**Authors:** Stephanie H Chen, Ashley Jones, Patricia Lu-Irving, Jia-Yee S Yap, Marlien van der Merwe, Jason G Bragg, Richard J Edwards

## Abstract

Technological advances have propelled DNA sequencing of non-model organisms, making sequencing more accessible and cost effective, which has also increased the availability of raw data in public repositories. However, contamination is a significant concern, and the use and reuse of sequencing data requires quality control and curation. A reference genome for the Australian native rainforest tree *Rhodamnia argentea* Benth. (malletwood) was assembled from Oxford Nanopore Technologies (ONT) long-reads, 10x Genomics Chromium linked-reads, and Hi-C data (N50 = 32.3 Mbp and BUSCO completeness 98.0%) with 99.0% of the 347 Mbp assembly anchored to 11 chromosomes (2*n* = 22). The *R*. *argentea* genome will inform conservation efforts for Myrtaceae species threatened by the global spread of the fungal disease myrtle rust. We observed contamination in the sequencing data and further investigation revealed an arthropod source. Here, we demonstrate the feasibility of assembling a high-quality gapless telomere-to-telomere mite genome using contaminated host plant sequencing data. The mite exhibits genome streamlining and has a 35 Mbp genome (68.6% BUSCO completeness) on two chromosomes, capped with a novel TTTGG telomere sequence. Phylogenomic analysis suggests that it is a previously unsequenced eriophyoid mite. Despite its unknown identity, this complete nuclear genome provides a valuable resource to investigate invertebrate genome reduction. This study emphasises the importance of checking sequencing data for contamination, especially when working with non-model organisms. It also enhances our understanding of two species, including a tree that faces substantial conservation challenges, contributing to broader biodiversity initiatives.

**Significance:** The genomes of *Rhodamnia argentea* and an associated eriophyoid mite, which contaminated the tree raw sequencing data, were assembled for the first time. We generated valuable chromosome-level genomic resources for the conservation of myrtle rust impacted tree species, pest genomics, and understanding genome streamlining. The research underscores the growing prevalence of sequencing experiments in non-model organisms while emphasising the importance of quality control and curation of sequencing data.

## Introduction

### Sequence contamination in the genomic era

The rapid development of sequencing technologies coupled with decreasing cost and have resulted in increased base accuracy, higher throughput, and longer read lengths (van Dijk et al. 2023); genomic resources are increasingly being generated for non-model species (Ekblom & Galindo 2011; Lewin et al. 2018). However, contamination in reference sequence databases and public sequence repositories is pervasive (Steinegger & Salzberg 2020; Lupo et al. 2021). Contamination during sample collection of sequence preparation can impact downstream analyses and interpretation (Laurin-Lemay et al. 2012) and studies have discovered cross-species contamination in genomes (Merchant et al. 2014).

The validation and curation of raw sequencing data and subsequent reference genome assembly is crucial for accurate and reliable genomes. Many tools for quality control and contaminant screens of sequencing data and assemblies are available (e.g. Andrews 2010; Kumar et al. 2013; Hadfield & Eldridge 2014; Cornet & Baurain 2022). The NCBI Foreign Contamination Screen (FCS) is a tool suite for identifying and removing contaminant sequences in genome assemblies. FSC was used to screen 1.6 million GenBank assemblies and identified 36.8 Gbp of contamination (0.16% of bases) (Astashyn et al. 2024). Screening vast volumes of data is intensive in multiple ways – computationally, financially, and time. The careful curation of a reference genome also requires these resources.

The use of genomics in plant biosecurity and conservation is growing, but requires molecular resources, such as reference genomes (Formenti et al. 2022; McMahon et al. 2014). Genomes provide a fundamental and comprehensive framework to understand genetic diversity and local adaptation. This can facilitate managers and stakeholders to make effective conservation decisions including quantifying genetic diversity and relatedness, understanding demographic history, and gene discovery (Fiedler et al. 2022; Rossetto et al. 2021; Supple & Shapiro 2018). Therefore, genomes are important for the conservation of species under threat from invasive species, including exotic plant diseases. For instance, the global spread of myrtle rust has become particularly concerning for many Australian native plants.

### Genomic resources for myrtle rust

Myrtle rust is caused by the fungal pathogen, *Austropuccinia psidii* (G.Winter) Beenken, and has a large global host list of 480 species in the family Myrtaceae that occur across a wide range of environments (Berthon et al. 2019; Soewarto et al. 2019). Myrtle rust originates from South America and was introduced to Australia in 2010. Since then, it has caused a rapid decline in susceptible native Myrtaceae species (Carnegie et al. 2016; Fensham & Radford-Smith 2021). An improved understanding of the genetic diversity and evolutionary history of impacted species will contribute to conservation management.

The Myrtaceae is a large family of woody plants containing over 5,650 species mainly occurring in the southern hemisphere (Govaerts 2008). There are several published Myrtaceae reference genomes, with sequencing efforts focused on improving breeding prospects for forestry and agriculture. *Eucalyptus grandis* W.Hill was the first published Myrtaceae reference genome (Myburg et al. 2014). Other commercially important species such as guava (*Psidium guajava* L.) (Feng et al. 2021) and clove (*Syzygium aromaticum* L.) (Ouadi et al. 2022) have also had their genomes published. Rapid technological advances in long-read sequencing present new opportunities for plant genomics (Dumschott et al. 2020; Xie et al. 2024). Notably, the chromosome-level gap-free T2T genome assembly of *Rhodomytus tormentosa* (Aiton) Hassk. using PacBio and Nanopore long-reads was published recently (Li et al. 2023). However, limited efforts have been directed towards generating genomic resources for Australian native plants impacted by myrtle rust, despite the value that genomes bring to conservation (Brandies et al. 2019). For example, there are currently no published *Rhodamnia* genomes, despite susceptibility of the genus to the disease (Carnegie et al. 2016; Pegg et al. 2014).

Trees act as hosts to a diverse range of pests and diseases (Gougherty & Davies 2022), and these present challenges for plant genome assembly. Many of these potential contaminants are too small to be observed with the naked eye and become inadvertently included in a sample. These pests and pathogens may get incorporated in plant genomes and issues can propagate as these genomic resources are reused. targeted sequencing of a host and/or its pest or pathogen may be complicated by these organisms all contaminating each other. For example, transcriptome sequencing of a predatory mite was contaminated by prey sequences due to an insufficient starvation period (Hoy et al. 2013).

### Advances in mite genomics

Eriophyoid mites (Acariformes: Eriophyoidea) are a hyperdiverse and widespread lineage of over 5,000 taxonomically accepted mite species that feed on plants (Zhang 2017; Ueckermann 2010). These four-legged mites have the distinct ability to induce galls in their plant hosts and many are agricultural pests (de Lillo et al. 2018; Lindquist et al. 1996). They are among the smallest of terrestrial arthropods, averaging 200 μm in length and difficult to visually detect (Amrine et al. 2003).

Progress in sequencing Trombidiformes, an order in the class Arachnida that includes eriophyoid mites, has accelerated in recent years, providing valuable evolutionary and applied insights. The two-spotted spider mite, *Tetranychus urticae* Koch, 1836, was the first complete chelicerate sequenced and was the smallest arthropod assembly at the time with a 90 Mbp genome (Grbić et al. 2011). This species is one of several agricultural pests sequenced and sheds light on pesticide resistance. Others include the false spider mite *Brevipalpus yothersi* Baker, 1949 (Navia et al. 2019), a fruit pest in Asia *Tetranychus piercei* McGregor, 1950 (Chen et al. 2024), the citrus red mite *Panonychus citri* (McGregor, 1916) (Yu et al. 2021), and the redlegged earth mite *Halotydeus destructor* (Tucker, 1925) which is invasive in Australia (Thia et al. 2023). Other mites such as *Pyemotes zhonghuajia* Yu, Zhang & He, 2010 have biocontrol potential (Song et al. 2022) while the mechanisms of gall formation have been studied through *Fragariocoptes setiger* (Nalepa, 1894) (Klimov et al. 2022). The size of the smallest sequenced Trombidiformes genome is 33 Mbp for the tomato russet mite *Aculops lycopersici* (Tryon, 1917) (Greenhalgh et al. 2020). On the other hand, the largest genome (193 Mbp) belongs to the chigger mite *Leptotrombidium pallidum* (Nagayo, Miyagawa, Mitamura & Tamiya, 1919) which is a major vector for scrub typhus (Kim et al. 2020). Other mites sequenced for human health include the European dust mite *Dermatophagoides pteronyssinus* (Trouessart, 1897) (Randall et al. 2018), *Dinothrombium tinctorium* (Linnaeus, 1767), and *Leptotrombidium delicense* (Walch, 1922) (Dong et al. 2018). Chromosome-level genomes are also emerging. For example, the genome of the free-living model species, *Archegozetes longisetosus* Aoki, 1965, provides insights into the evolution of parasitic lifestyles (Brückner et al. 2022).

### Aims of the study

Here, we present a chromosome-level reference genome of *Rhodamnia argentea* Benth. (malletwood) which is variably resistant to myrtle rust (Pegg et al. 2014) assembled from Oxford Nanopore Technologies (ONT) long-reads, 10x Genomics Chromium linked-reads, and Hi-C data. We highlight the importance of checking for contamination in genome sequencing projects, and provide an example of this decontamination process, through the long-read assembly of a previously unsequenced eriophyoid mite. We analyse the mite genome and provide insights into genome streamlining and a novel telomere sequence.

## Materials and methods

### Sampling and sequencing

Young leaves of *Rhodamnia argentea* from a tree at the Royal Botanic Garden Sydney (NCBI BioSample SAMN19698777, ToLID drRhoArge1) were sampled and kept moist in a zip lock bag and refrigerated at 4°C until extraction. Additionally, 5 g of young leaf tissue was snap frozen in liquid nitrogen immediately after collection for Hi-C.

For the ONT long-read sequencing, HMW gDNA was obtained using two sorbitol pre-washes (Inglis et al. 2018) followed by extraction using a magnetic bead-based protocol described in Jones et al. 2021. A clean-up and size selection step were performed to remove DNA fragments <40 kb using the BluePippin 0.75% Agarose Gel Cassette, Dye Free on the BluePippin (Sage Science, Beverly, MA, USA).

High-molecular-weight gDNA was sent to AGRF for 10x Genomics Chromium sequencing. Size selection was performed to remove DNA fragments <40 kb using the BluePippin 0.75% Agarose Gel Cassette, Dye Free on the BluePippin (Sage Science, Beverly, MA, USA). Briefly, 5 µg of DNA was diluted to 30 µL in TE buffer and 10 µL of room temperature equilibrated loading buffer was added to each aliquot and mixed by pipetting. Samples were loaded on the cassette by removing 40 µL of buffer from each well and adding 40 µL of sample or external marker. The cassette was run with the 0.75% DF Marker U1 high-pass 30-40 kb v3 cassette definition. Size selected fractions (approximately 40 µL) were collected following the 30 min electrophoresis run. The library was prepared using the Chromium Genome Library Kit & Gel Bead Kit according to the manufacturers’ instructions. Sequencing was performed on an Illumina NovaSeq 6000 with a SP flow cell at 300 cycles (2 x 150 bp paired-end sequencing) using an XP 2-Lane Kit for individual lane loading.

Sequencing was performed on the MinION Mk1B using a FLO-MINSP6 (R9.4.1) flow cell with a library prepared with the native DNA ligation kit (SQK-LSK109). Basecalling was performed after sequencing with GPU-enabled Guppy v3.2.1. The output from all ONT basecalling was pooled for adapter removal using Porechop v.0.2.4 (Wick et al. 2017) and quality filtering (removal of reads less than 500 bp in length and Q lower than 7) with NanoFilt v2.6.0 (De Coster et al. 2018) followed by assessment using FastQC v0.11.8 (Andrews 2010).

For the 10x linked-read sequencing, high-molecular-weight (HMW) genomic DNA (gDNA) was obtained using a sorbitol pre-wash step prior to a CTAB extraction adapted from Inglis et al. (2018). The gDNA was then purified with AMPure XP beads (Beckman Coulter, Brea, CA, USA) using a protocol based on (Schalamun et al. 2019) (Lu-Irving & Rutherford 2021). The quality of the DNA was assessed using Qubit, NanoDrop and TapeStation 2200 System (Agilent, Santa Clara, CA, USA).

Hi-C library preparation and sequencing for *R. argentea* leaf tissue was conducted at the Ramaciotti Centre for Genomics, using the Phase Genomics Proximo Hi-C Plant kit using Sau3AI to digest the genome. The library was assessed using Qubit and the Agilent 2200 TapeStation system (Agilent Technologies, Mulgrave, VIC, Australia). A pilot run on an Illumina iSeq 100 with i1 300 cycle chemistry (2 x 150 bp paired end sequencing) was performed, for QC using hic_qc v1.0 (Phase Genomics 2019). This was followed by sequencing on the Illumina NextSeq 500 using a High Output 300 cycle kit, v2.5 chemistry (2 x 150 bp paired-end sequencing).

### Genome assembly and decontamination

A draft genome assembly for *R. argentea* was assembled from the 10x linked-read sequencing using Supernova v2.1.1 (Weisenfeld et al. 2017). Following an initial assembly using all the data, a second assembly was generated subsampling to 300 million reads and 50% barcode retention (--maxreads=300000000 --bcfrac 0.5) to account for high DNA concentration and a small genome. This resulted in a raw coverage of 54.93X (ideal ∼56X) and effective coverage of 42.95X (ideal ∼42X) as reported by Supernova. Supernova ‘pseudohap2’ was converted into non-redundant primary and alternative assemblies using custom code now available as part of Diploidocus (runmode=dipnr) (Stuart et al. 2022).

Following the release of the unpublished draft assembly (v1) on NCBI (GCF_900635035.1), possible contamination was identified (see Discussion). We therefore generated additional ONT MinION and Hi-C data reassembled the *R. argentea* v2 genome with a targeted contaminant removal step, before assembling contaminant reads into the mite genome (Figure 1). *Rhodamnia argentea* ONT MinION reads were assembled using NECAT (Chen et al. 2021) v0.01 followed by tidying with Diploidocus v0.17.0 (Chen et al. 2022) on cycle mode along with contaminant screening and purging of the myrtle rust pathogen genome (Edwards et al. 2022). Convergence was reached in four rounds. The assembly was long-read polished with Racon v1.4.5 (Vaser et al. 2017) using the parameters -m 8 -x -6 -g -8 -w 500 and medaka v1.0.2 using the r941_min_high model. Then, the 10x reads were incorporated by short-read polishing using Pilon v1.23 (Walker et al. 2014) with reads mapped using Minimap2 v2.12 (Li 2018) and correcting for SNPs and indels. Scaffolding was done using SSPACE-LongRead v1.1 (Boetzer & Pirovano 2014) followed by gap-filling using gapFinisher v20190917 (Kammonen et al. 2019) with default parameters. After another round of long-read polishing with Racon v1.4.5 (Vaser et al. 2017) and medaka v1.0.2, we moved forward with a second round of tidying in Diploidocus (default mode).

**Figure 1.**
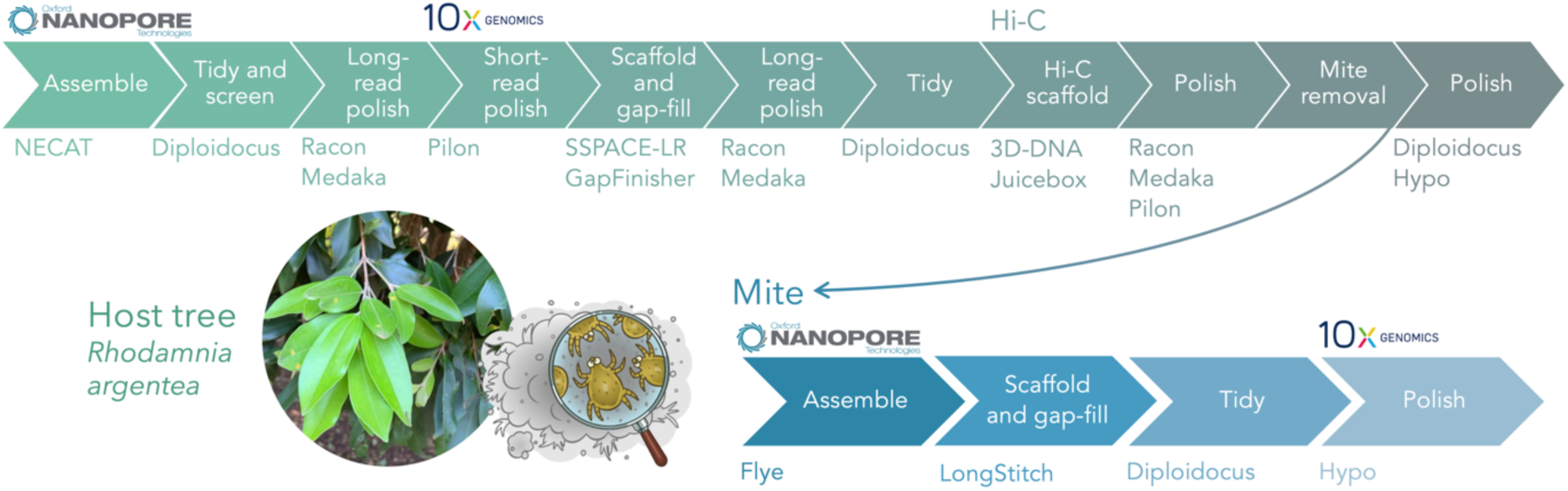
Assembly workflow of the host tree, *Rhodamnia argentea*, and associated mite genome. Tree photo by SH Chen.

The Hi-C reads were aligned to the draft genome assembly using the Juicer pipeline v1.6 (Durand et al. 2016) then scaffolds were ordered and orientated using the 3D *de novo* assembly pipeline v180922 (Dudchenko et al. 2017). The contact map was visualised using Juicebox Assembly Tools v1.11.08 and errors over 2 review rounds were corrected manually to resolve 11 chromosomes (Dudchenko et al. 2018). Additionally, the individual’s diploid status was confirmed using Smudgeplot v0.2.1 (Ranallo-Benavidez et al. 2019). Then followed another round of long-read polishing with Racon v1.4.5 (Vaser et al. 2017) and medaka v1.0.2, and tidying in Diploidocus (default mode). A final polish was performed using Hypo v1.0.3 as it improved the base accuracy of the assembly (Kundu et al. 2019).

We are also working on a critically endangered species of *Rhodamnia*, *R*. *rubescens*, and mapped the *R*. *argentea* scaffolds to the unpublished *R*. *rubescens* genome; the *R. rubescens* DNA was extracted from tissue culture i.e. sterile and contaminant free. Taxolotl v0.1.3 (Tobias et al. 2022) was used to investigate the taxonomy of the mapped and unmapped sets of scaffolds. Briefly, Taxolotl uses mmseqs2 (Steinegger & Söding 2017) to search protein sequences against the NCBI nr database (compiled 09/03/2021) and assign taxonomy before mapping them back onto contigs and scaffolds. In addition to generating results for all annotation proteins, taxonomic assignments are made for each scaffold, which are rated as ‘good’ or ‘bad’ based on the dominant taxon. Results can then be visualised with Pavian v1.2.0 (Breitwieser & Salzberg 2020). In this case, because the draft *R. argentea* genome was already in NCBI, the nr database used for Taxolotl had *R. argentea* screened out.

All scaffolds appeared to be either pure tree or pure contamination; the 74 mapped scaffolds (347 Mbp) were 99.93% plant and 12 scaffolds that failed to map (34.6 Mbp) were 97.8% animal (Supplementary Figure 4). The latter set of scaffolds were removed from the *R. argentea* genome (except for Rarg_v2.81 which was deemed to be of plant origin after additional Taxolotl analysis and phylogenetic analysis) and a final polish was performed using Hypo v1.0.3 for the final Rarg_v2 assembly (drRhoArge1.2).

ONT and 10x reads were partitioned into three sets based on mapping to the contaminated v2 genome: (1) Rarg_v2 reads, (2) contamination reads mapping to removed scaffolds, and (3) unmapped reads. Contamination and unmapped reads were then combined for each technology, with 10x reads trimmed with bbmap v38.51 (Bushnell 2014) (5’ 30 bp of R1, 5’ 10 bp of R2, quality trimming <Q20 from both ends) and filtered to keep read pairs where both sequences were at least 100 bp long. Non-Rarg_v2 ONT reads were assembled with Flye v2.9 (Kolmogorov et al. 2019) with a target genome size (based on the filtered scaffolds) of 35 Mbp. Scaffolding and gap-filling was done with LongStitch-ARKS v1.0.1 (Coombe et al. 2021). A tidy was done with Diploidocus on cycle mode (Chen et al. 2022). The genome was then polished with Hypo v1.0.3 (Kundu et al. 2019) using partitioned 10x reads as short-read data, mapped with BWA v0.7.17 (Li 2013), and ONT reads as long-read data, mapped with minimap2 v2.22 (Li 2018).

Both the *R*. *argentea* and mite assemblies were subject to a final contamination screen by running FCS-GX v0.4.0 (Astashyn et al. 2024) and Tiara v1.0.3 (Karlicki et al. 2022). FCS-GX was run against the NCBI gxdb (build date 2023-01-24; downloaded 2023-09-12). DepthKopy analysis of the mite assembly was also performed to check for low quality/coverage sequences; plots were generated with ggstatsplot (Patil 2021) with box plots marking the median and quartiles.

### Assembly completeness and genome size

Assembly completeness of the *R*. *argentea* genome was evaluated by with BUSCO v5.3.0 (Manni et al. 2021) against the eudicot_odb10 dataset (*n* = 2,326) using MetaEuk (Levy Karin et al. 2020) as the gene predictor. For the mite genome, we ran BUSCO v5.2.2 against the arachnida_odb10 (*n* = 2,934), arthropoda_odb10 (*n* = 1,013), metazoa_odb10 (*n* = 954), eukaryota_odb10 (*n* = 255) datasets using MetaEuk as the gene predictor. DepthSizer v1.6.2 (Chen et al. 2022) was used for genome size estimation from the read depth of single-copy orthologues for the *R*. *argentea* genome and to investigate the read depth of the mite genome contamination.

### Genome annotation

The *R. argentea* genome was annotated using the homology-based gene prediction program GeMoMa v1.7.1 (Keilwagen et al. 2019) with four reference genomes downloaded from NCBI: *Eucalyptus grandis* (GCF_000612305.1) (Myburg et al. 2014), *Syzygium oleosum* (v1 10x linked-read genome; GCF_900635055.1) (Edwards, unpublished), *Rhodamnia argentea* (v1 10x linked-read genome; GCF_900635035.1) (Edwards, unpublished), and *Arabidopsis thaliana* (TAIR10.1, GCA_000001735.2) (Swarbreck et al. 2007). Annotation completeness was assessed using BUSCO in proteome mode. Ribosomal RNA (rRNA) genes were predicted with Barrnap v0.9 (Seemann 2018) and transfer RNAs (tRNAs) were predicted with tRNAscan-SE v2.05 (Lowe & Chan 2016), implementing Infernal v1.1.2 (Nawrocki & Eddy 2013) and recommended filtering for eukaryotes for form the high-confidence set.

The mite genome was also annotated with GeMoMa v1.7.1 with 15 invertebrate genomes annotated by NCBI (Supplementary Table 1). Predicted transcriptome statistics were generated with SAAGA v0.7.7 (Edwards et al. 2021).

For both genomes, a custom repeat library was generated with RepeatModeler v2.0.1 (-engine ncbi) and the genome was masked with RepeatMasker v4.1.0 (Tarailo-Graovac & Chen 2009), both with default parameters. The annotation table was generated using the buildSummary.pl RepeatMasker script. Telomeres repeat sequences were identified by manually inspecting the ends of the two chromosome-scale scaffolds in Mite_v1 and creating a consensus repeat unit.

### Mite phylogenomics and comparative genomics

To put the sequenced mite genome in context, 36 mite genomes were downloaded from NCBI [04 Nov 2021] (Supplementary Table 1) and subject to the same BUSCO, GeMoMa, SAAGA and RepeatModeler/RepeatMasker annotation as our mite genome (hereafter, Mite_v1). BUSCO results for all the mite species and selected taxonomic levels were compiled using BUSCOMP v1.1.3. For each taxonomic grouping of species, the ‘best’ rating for each gene was retained, where Single > Duplicated > Fragmented > Missing.

To identify the sequenced mite, phylogenomic analysis was done with the BUSCO single copy orthologues from the arachnida_odb10. To avoid biases due to genome reduction, only Complete genes from Mite_v1 were used for the analysis. For each BUSCO gene, HAQESAC v1.14.0 (Edwards et al. 2007) was used to generate protein multiple sequence alignments and phylogenetic trees for all species returning a single copy orthologue for that gene. Briefly, HAQESAC compares all proteins to the Mite_v1 query and removes sequences too distant or incomplete, before aligning with Clustal Omega 1.2.4 (Sievers et al. 2011). Poorly aligned sequences were removed by HAQESAC and a phylogeny inferred by FastTree v2.1.11 (Price et al. 2010) before being mid-point rooted. Trees were then combined into a consensus tree with ASTRAL v5.7.8 (Zhang et al. 2018). Synteny analysis based on BUSCO genes (arachnida_odb10 database) was used to compare the mite genome to ACULYCv1 (Greenhalgh et al. 2020) and visualised with ChromSyn (Edwards et al. 2022). Telomere prediction was performed using Telociraptor v0.9.0 (Edwards 2023) with forward and reverse regular expressions C{1,3}A{2,4} and T{2,4}G{1,3}, based on the TTTGG telomere repeat identified. Additional telomeric repeats were identified using TIDK v0.2.31 (Brown et al. 2023) search with the telomere sequence CCAAA. Ribosomal rDNA genes were predicted using barrnap v0.9.0 (Seemann 2018) in eukaryotic mode.

## Results

Nanopore MinION sequencing yielded 21.7 Gbp whilst 10x linked-read sequencing yielded 140 Gbp for *Rhodamnia argentea* (Supplementary Table 2). The genome size of the studied species had not been previously measured using flow cytometry and the assembly size was 347 Mbp and DepthSizer predicted, on average, 365 Mbp (Supplementary Table 3). The chromosome-level genome had a high structural completeness with a scaffold N50 of 32.3 Mbp.

The diploid status of the *R*. *argentea* individual was confirmed using Smudgeplot (Supplementary Figure 1) and 11 chromosomes were resolved, with unplaced scaffolds (Supplementary Figure 2). The anchored portion of the genome was 343,265,630 bp, accounting for 99.0% of the assembly. Rarg_v2 comprised a total of 75 scaffolds (Table 1). Comparing the v2 genome to the v1 genome assembled using only the 10x linked-reads, the contiguity of the v2 genome is vastly improved. The Rarg_v1 genome (GCF_900635035.1) consists of 15,781 unplaced scaffolds (N50 of 857 kb) and 23,310 contigs (N50 of 63.7 kb). However, the BUSCO completeness is comparable to v2 at 98%. There was a low number of duplicated single copy BUSCOs overall. The 34.5 Mbp mite genome consisted of two gapless telomere-to-telomere chromosomes and five additional contigs on four scaffolds (Table 1).

**Table 1.** Genome assembly and annotation statistics.

| Statistic | Rarg_v2 | Mite_v1.1 |
| --- | --- | --- |
| Technology | ONT (63x), 10x (404x), Hi-C | ONT, 10x |
| **Total length (bp)** | **346,714,100** | **34,344,194** |
| **No. of scaffolds** | **75** | **2** |
| N50 (bp) | 32,337,726 | 17,366,404 |
| L50 | 5 | 1 |
| **No. of contigs** | **111** | **2** |
| N50 (bp) | 14,585,590 | 17,366,404 |
| L50 | 10 | 1 |
| N bases | 3,600 | 0 |
| GC (%) | 40.38 | 42.08 |
| **BUSCO complete (genome)** | **98.0% (2,280)** | **68.6% (2,012)** |
| Single-copy (genome) | 94.7% (2,203) | 64.1% (1,880) |
| Duplicated (genome) | 3.3% (77) | 4.5% (132) |
| BUSCO fragmented (genome) | 0.7% (17) | 1.3% (38) |
| BUSCO missing (genome) | 1.3% (29) | 30.1% (883) |
| **Protein-coding genes** | **35,646** | **8,050** |
| mRNAs | 51,034 | 9,244 |
| rRNAs | 697 | 18 |
| tRNAs | 408 | 56 |
| **BUSCO complete (proteome)** | **96.0% (1,382)** | **70.3% (2,063)** |
| Single-copy (proteome) | 76.5% (1,101) | 65.1% (1,909) |
| Duplicated (proteome) | 19.5% (281) | 5.2% (154) |
| BUSCO fragmented (proteome) | 1.2% (18) | 1.3% (38) |
| BUSCO missing (proteome) | 2.8% (40) | 28.4% (833) |

BUSCO completeness (arachnida_odb10, *n =* 2,934) was low, at 68.6%, with a surprisingly high Duplicated rate of 4.6%. Whilst 30.1% BUSCO genes are missing, only 1.3% were fragmented, which is comparable to the Rarg_v1 genome. DepthKopy predicted that 129/135 (95.6%) of the Duplicated genes are real duplications, consistent with the gapless telomere-to-telomere genome of high quality. BUSCO completeness broadly increased as the taxonomic specificity was reduced (Supplementary Table 4), with 87.4% eukaryotic genes (*n* = 255) Complete and only 6.7% Missing.

Genome annotation predicted 35,646 protein-coding genes and 51,034 mRNAs, along with 697 rRNAs and 408 tRNAs. The BUSCO completeness for the proteome was 96.0% (Table 1). In contrast, the GeMoMa annotation of the mite genome predicted only 8,071 protein-coding genes (9,269 mRNAs), with 18 rRNAs and 56 tRNAs. The 18 rRNA genes comprised six tandem copies of the complete 18S-5.8S-28S rDNA repeat unit. DepthKopy analysis (Figure 2) revealed that the rDNA cluster was collapsed in the assembly and the six copies of the repeat unit represented 36 copies in the actual genome (mean CN per gene ∼12).

**Figure 2.**
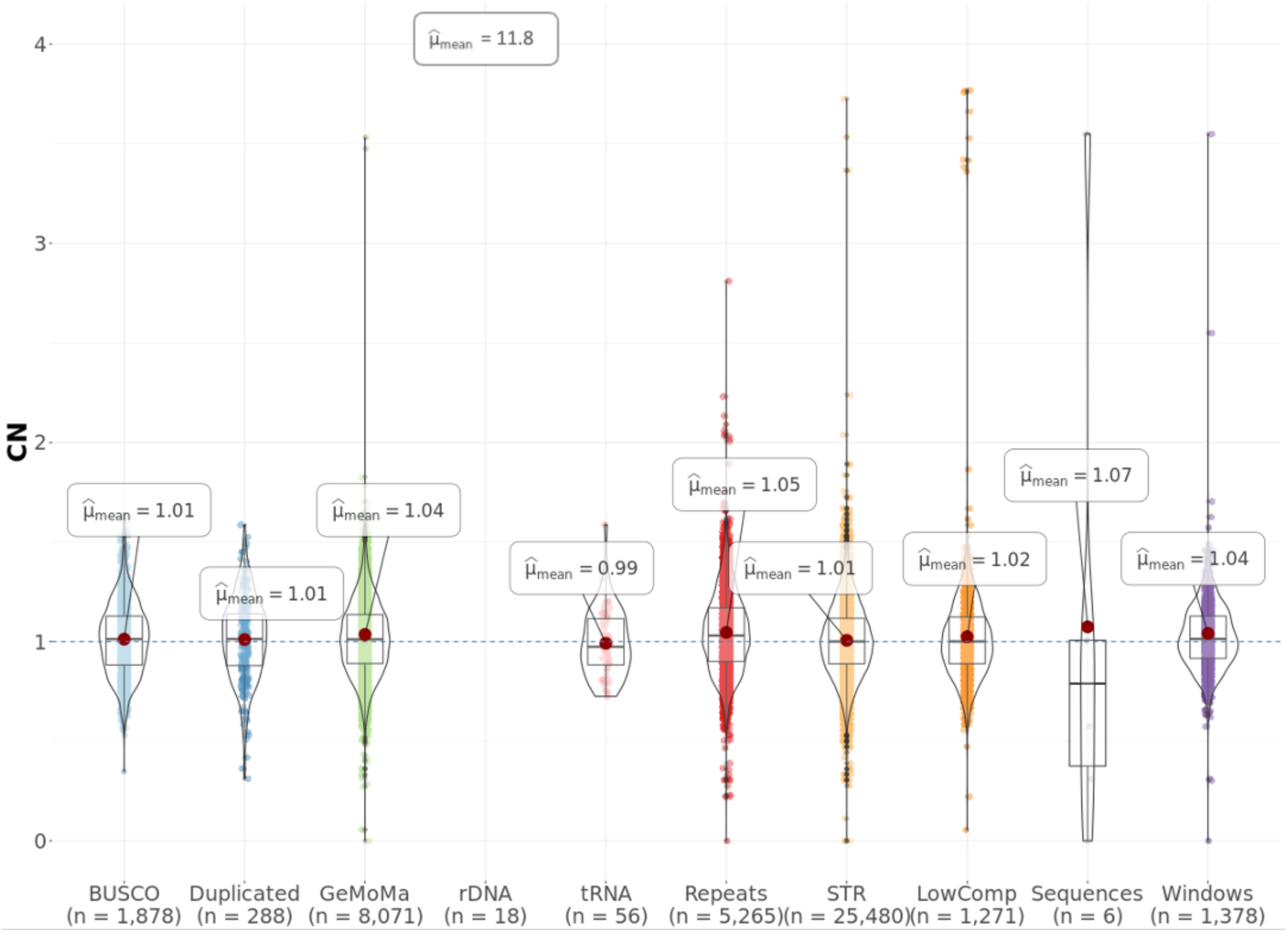
DepthKopy predicted copy number distribution for the assembled mite. Mapped ONT read depths have been converted into copy number distributions using the modal single-copy read depth from ‘Complete’ BUSCO genes of 35.7X. Whiskers extend to the most extreme values no further than 1.5 times the inter-quartile range. BUSCO, single-copy complete BUSCO genes; Duplicated, duplicated complete BUSCO genes; GeMoMa, GeMoMa gene models; rDNA, rDNA gene models; tRNA, tRNA gene models; Repeats, complex repeats; STR, short tandem repeats; LowComp, low complexity regions; Sequences, individual scaffolds; Windows: 25 kb tiled windows.

The *R. argentea* genome is highly repetitive with a repeat content of 43.6% (Supplementary Table 5). The mite genome had a low repeat content at 11.1% (Supplementary Table 6). There was no sign of widespread collapsed repeats in the assembly, with most repeats having a consistent copy number around 1 (Figure 2).

The *R*. *argentea* reads had a mean read depth of approximately 53X whilst the contaminants were at 37X (Supplementary Figure 3). The contaminating organisms were not obvious on the leaf sample from examination with the naked eye at the time of sample collection and wet lab work. Nor was it obvious what the contaminating organism looked like during subsequent examination using a dissecting microscope after sequencing was completed.

Taxolotl identified the 74 scaffolds that were mapped to the *R. rubecens* genome as plant and 12 unmapped scaffolds as primarily Trombidiforme mites (Supplementary Figure 4). Of the six scaffolds reassembled from the contaminated and unmapped reads, two were gapless telomere-to-telomere chromosomes that comprised 99.64% of the assembly. These were capped with telomeres that had a novel TTTGG sequence.

After quality and contamination assessment, the four unplaced scaffolds totalling 124,586 bp were removed as potential contamination (1 scaffold, 2 contigs) or a false duplication (1 contig). The size of the assembled mite genome is comparable to two previously published eriophyid mite genomes: *Aculops lycopersici* (Greenhalgh et al. 2020) and *Fragariocoptes setiger* (Klimov et al. 2022). Phylogenomic analysis of BUSCO genes with 36 mite genomes from NCBI (Supplementary Table 1) confirmed that the assembled mite was within the Trombidiformes, in the superfamily Eriophyoidea (Figure 3a), with the closest available genome from *Aculops lycopersici* (Greenhalgh et al. 2020) in the family Eriophyidae.

**Figure 3.**
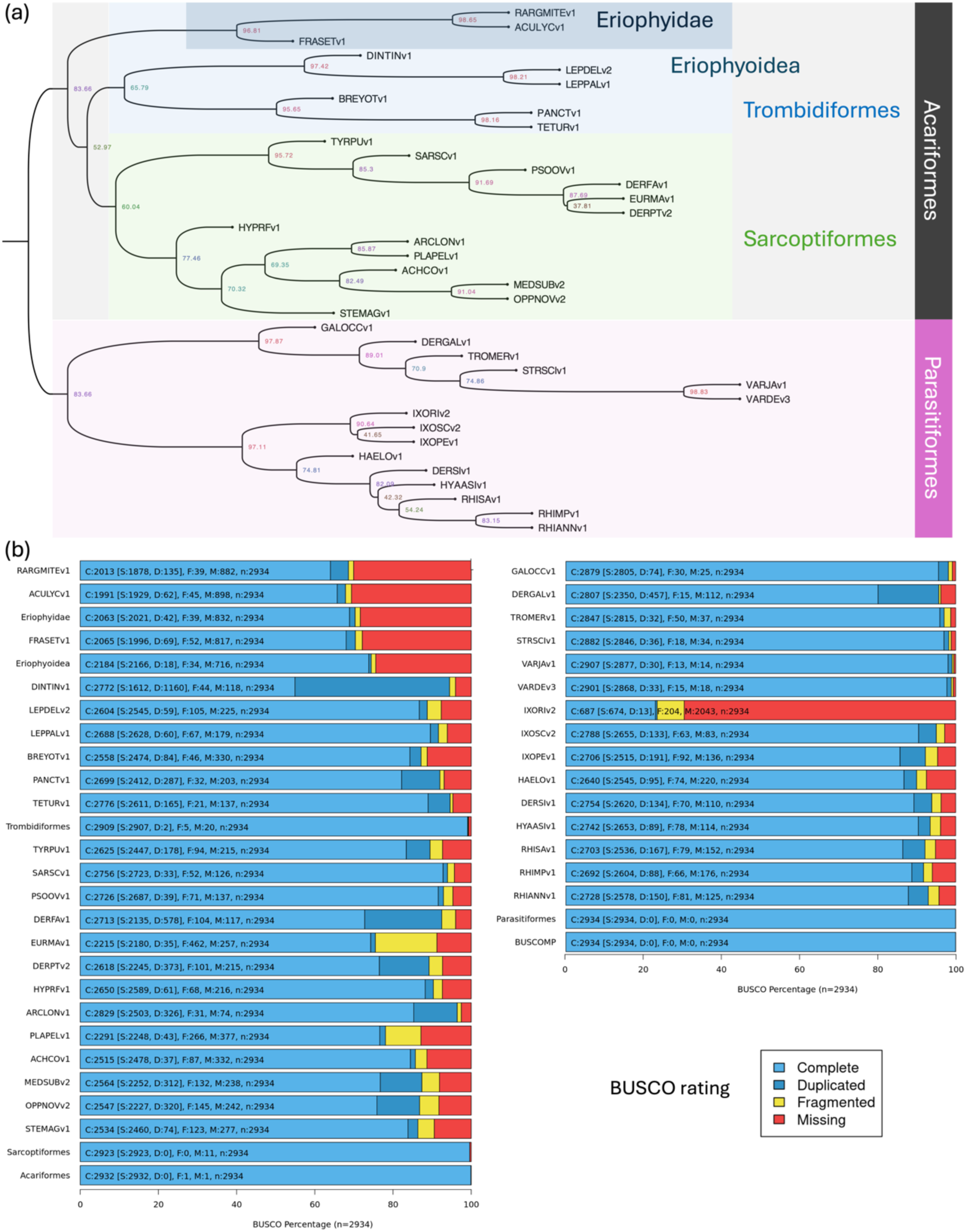
Mite phylogenomics and genome reduction. (a) ASTRAL consensus tree for 1,877 BUSCO Complete gene trees (arachnida_odb10). Internal nodes are labelled with the percentage of quartets in the BUSCO gene trees that support the ancestral branch. Select taxonomic groupings are marked (see text for details). Full species names for each assembly can be found in Supplementary Table 1. (b) BUSCO completeness for each assembly and BUSCOMP compilation for taxonomic groups. For each taxonomic grouping in the phylogenomic tree, numbers report the best ratings for each BUSCO gene.

Examining synteny of the assembled mite to the tomato russet mite (*Aculops lycopersici*) via Complete BUSCO genes, there are extensive rearrangements with 71% protein sequence identity (Figure 4). Both Eriophyoidea species showed similarly low BUSCO completeness with many of the same genes Missing. Of the 882 BUSCO genes missing in our mite genome, 832 were also missing in *A. lycopersici* (Figure 3b, Eriophyidae) and 716 of these were missing in *Fragariocoptes setiger* (Figure 3b, Eriophyoidea). In contrast, only 20 genes (0.7%) were also missing across all remaining Trombidiformes. The low BUSCO completeness and high missing values in these three genomes are consistent with genome reduction when compared to the other 34 mite genomes analysed. These species also show a reduction in the number of introns and transposable elements, and an overall reduction in repetitive and intergenic content (Supplementary Figure 5). Despite a reduction in gene numbers, the percentage of the genome that corresponds to exons and genes is higher in these species (Supplementary Figure 5a).

**Figure 4.**
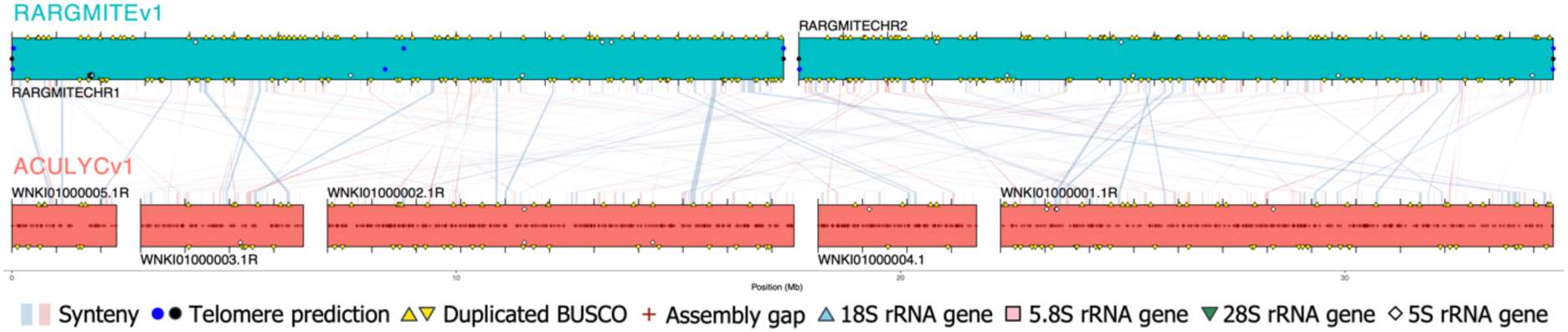
Synteny between assembled mite and *Aculops lycopersici* (tomato russet mite). Coloured boxes represent scaffolds containing Complete BUSCO genes. Blue and red blocks represent blocks of collinear and inverted synteny based on 2+ adjacent BUSCO genes in the same relative orientation. Black circles, Telociraptor telomere predictions; Blue circles, TIDK-predicted telomere repeats; Red pluses, assembly gaps; Yellow triangles, Duplicated BUSCO genes; Blue triangle, 18S rRNA gene; Pink square, 5.8S rRNA gene; Green triangle, 28S rRNA; White diamond, 5S rRNA gene.

## Discussion

### Identification and removal of assembly contamination

Improvements in sequencing technology over recent years have massively increased the ease with which whole genomes can be sequenced and assembled (Gupta 2022). However, as the scale of genomes assembled rise, the relative resources available per genome for curation decreases (Howe et al. 2021). This can be particularly challenging when project priorities lie elsewhere. Genome contamination is a well-known problem and is particularly prevalent among short-read and unpublished draft genomes (Kumar et al. 2013; Chrisman et al. 2022; Steinegger & Salzberg 2020). The mite contamination in the *R. argentea* draft genome was particularly difficult to identify in the initial assembly due to the large number of variable-quality contigs, the high depth of contaminant sequencing, and the lack of genome assemblies from closely related organisms. Variations in read depth and GC content were discounted as possible regional biases within assembled scaffolds, such as repetitive regions with differences in copy number between the assembly and the true genome. Instead, contamination in this case was identified by targeted searches for arthropod sequences (pers. comm. W. Dermauw, J. Santolini, and S. Gerber).

More recently, numerous tools have become available to improve the ease and resolution of genomic DNA contamination screening, including Blobtoolkit (Kumar et al. 2013), FCS-GX (Astashyn et al. 2024), Conterminator (Steinegger & Salzberg 2020), Tiara (Karlicki et al. 2022), and ContScout (Bálint et al. 2024). The approach used here, Taxolotl (Tobias et al. 2022), complements these DNA-based methods by assigning taxonomy to protein sequences, which can enable matches across greater evolutionary distances. The exponential availability of genome assemblies across taxa, combined with the routine FCS-GX screening by NCBI upon submission, should greatly reduce the number of new genomes with significant contamination. Nevertheless, caution should always be used when interpreting unusual findings (e.g. Saffar & Matin 2021). This is particularly true when using unpublished draft genomes, where it is always good practice to contact the submitting researchers and discuss the quality of the assembly. Given the ongoing advances in both the methods and data required to detect contamination (namely improved genomic resources for possible contaminants), it would be a sensible precaution to run FCS-GX and/or Tiara on downloaded genomes prior to use where contamination might lead to misleading results. Long-term, the community needs improved mechanisms by which contamination in genomes can be removed from public repositories. Ideally, expert institutions (e.g. zoological/botanical gardens and museums) will be given the resources and power to leverage high-confidence data and taxonomic knowledge to identify and flag contamination for removal.

### Extraction and sequencing choices can reduce contamination risks

Improvements in sequencing and assembly methods also have an important role to play in reducing contamination. Long-read technologies are less prone to misincorporation of contamination during assembly. The quality of the DNA is a major determinant of sequencing success (Schalamun et al. 2019). Characteristically, long-read technologies require clean high quality HMW DNA, with the length and integrity of molecules being particularly important. Obtaining high quality DNA can be challenging for non-model organisms where extraction protocols have not been previously tested nor optimised; it is recommended that multiple extraction protocols are tested before sequencing a new plant species (Dumschott et al. 2020). For example, Myrtaceae contain many secondary compounds such as terpenes (Retamales & Scharaschkin 2014; Keszei et al. 2008) that can be problematic for sequencing. Techniques such as sorbitol washes, nuclei preparation, size selection and magnetic bead clean-up can lead to better quality DNA. Further, often sequencing centres will perform additional clean-up steps before library preparation.

Open science, including researchers sharing knowledge on platforms such as the Oxford Nanopore community forums and protocols.io (Teytelman et al. 2016), will facilitate successful DNA extraction and avoid constant ‘reinvention of the wheel’ (Schalamun et al. 2019). The sharing of expertise and experience of researchers optimising HMW DNA extraction specifically for long-read sequencing for species including Myrtaceae species such as *Eucalyptus* (Jones et al. 2021) provided benefits to this project. *Rhodamnia rubescens* (Benth.) Miq. tissue culture was available as the species is being conserved ex situ due to myrtle rust (Sommerville et al. 2019). The sequencing data generated from that species was used to decontaminate the *R. argentea* genome due to the sterile nature and lack of secondary compounds of tissue culture. Therefore, we recommend DNA extraction from tissue culture where it is available for a study species.

### A gapless telomere-to-telomere reference genome for an unidentified Eriophyoid mite

Whilst the relatively high abundance of the contaminating mite DNA made it challenging to identify in the draft *R. argentea* genome, the subsequent availability of ONT reads enabled the first gapless telomere-to-telomere assembly of a mite genome. Both chromosomes were capped with clearly identifiable telomeric repeats. The novel TTTGG sequence identified could improve telomere identification in other mites if this is a shared trait. Phylogenomic analysis was able to position the new mite within Eriophyoidae, but even the closest sequenced relative – the tomato russet mite, *Aculops lycopersici* (Greenhalgh et al. 2020) – was quite distantly related, with little conserved synteny (Figure 4) and a mean BUSCO protein sequence identify of 71%.

Attempts to physically identify the contaminating mite were not successful. It was not obvious what the contaminant was from observing leaves from the reference genome tree in the Royal Botanic Garden Sydney with the naked eye nor a dissecting microscope. The fact that the mite was collected and sequenced with the tree on separate sampling events years apart (first for 10x linked-reads then ONT and Hi-C), means that the contaminant was likely present in abundance and not removed by rinsing the leaves in water. Eriophyoid mites are microscopic and can reside within host tissue, presenting additional challenges for identification and removal (Amrine et al. 2003; Amrine & Manson 1996).

The Eriophyoidea is the most species-rich superfamily of the mites (Acari) and are found across the globe. They are all phytophagous and more than 80% of the group have high host specificity (Skoracka et al. 2010). While we do not have any information on the mite taxonomy or host specificity of the sequence we report on, Scott (1978) recorded that many herbarium specimens of *R*. *argentea* had densely tomentose fruits that were deformed by galls that could be mite induced. *Rhodamnia argentea* also has hairy lower leaf surfaces providing suitable habitat for phytophagous mites. However, vagrant mite species (non-gall forming) are the less host specialised and occur within this superfamily and can be found on multiple families or species (Oldfield 1996).

Eriophyoid mites can serve as vectors for viruses (de Lillo et al. 2018) and have also been observed as vectors for the rust fungus *Puccinia* spp. (Gamliel-Atinsky et al. 2010). Given the severity of the threat of myrtle rust to species of *Rhodamnia*, future work may include more detailed investigations into these mites particularly into the potential role as vectors. While the relationship between mites and their host plants were not the subject of our investigations, we show here how utilising ‘contamination’ can help further gain knowledge of the largely unknown and unexplored world of mite and plant host interactions. Presence of Eriophyoid mites can be seasonal and these tiny organisms (averaging approximately 200–300 µm) are not always detectable (Chetverikov et al. 2022) however we will continue to make observation throughout the year to further advance knowledge of the organism behind the genome.

At 32.5 Mbp, the *A. lycopersici* genome is the smallest yet reported for any arthropod and, reminiscent of microbial eukaryotes, exceptionally streamlined. It has few transposable elements, tiny intergenic regions, and is remarkably intron-poor, as more than 80% of coding genes are intronless (Greenhalgh et al. 2020). At 34.3 Mbp, our Rarg_mite_v1 assembly is not much bigger, and is notable in its completeness despite apparently poor BUSCO results. The latter observation appears to be the consequence of genome reduction, in which many expected genes have been lost (Arkhipova 2018; Hessen et al. 2010). Indeed, most non-tick mites have small genomes (Gregory & Young 2020). The reasons for the disparity in genome size between ticks and other mites remain unclear, but it is plausible that the small body size of the mites are made possible by streamlined genomes and/or that the evolution of small bodies has constrained genome size compared to other arthropods (Gregory & Young 2020). Genome streamlining is a fascinating biological phenomenon and increased genomic mite resources will broaden our understanding of the evolutionary mechanisms and genetic underpinnings responsible for reduced genome sizes. Most of the missing BUSCO genes are shared between Eriophyoidea but not other Trombodiform mites (Figure 3b), and presumably represent initial streamlining that occurred early in this lineage. However, continued lineage-specific losses suggest that genome reduction is an ongoing process. This is further supported by a surprisingly high Duplicated rate, which appear to represent genuine duplications within the mite lineage, consistent with ongoing adaptive evolution. It is also notable that the proportion of missing BUSCO genes increased considerably with increasingly taxonomically-restricted BUSCO datasets, suggesting that it is more derived traits that are being altered.

### Expanding genomic resources for Myrtaceae conservation and research

The *R. argentea* reference genome represents a genomic resource that will facilitate further efforts in Myrtaceae conservation, management, and research. Given that Myrtaceae is a large and diverse plant family with many economically important species, genomic resources are continually being generated in this family — a reflection of improved sequencing technologies, bioinformatic tools and knowledge. We provide the first *Rhodamnia* genome, a diverged Myrtaceae genus of rainforest trees. The genome fills a taxonomic gap and serve as reference point for species classification and comparative genomics in Myrtaceae. Current Myrtaceae research has focused on *Eucalyptus* (Ferguson et al. 2024), *Corymbia* (Healey et al. 2021), *Melaleuca* (Chen et al. 2023), *Syzygium* (Ouadi et al. 2022, 2023), and *Rhodomyrtus* (Li et al. 2023). These genomes have differing sizes of approximately 300–600 Mbp yet amazingly retain a conserved karyotype. However, many Myrtaceae species are under threat due to myrtle rust. Further research is needed to understand resistance to myrtle rust, and potential genomic ‘arms race’ with the causal agent’s much larger 1 Gbp genome (Tobias et al. 2021).

### Future directions in sequencing host plant with pests and pathogens

Another opportunity in the field of pest and pathogen genomes is an integrative approach where the host and symbiont are sequenced and assembled simultaneously as a metagenome. This will accelerate our knowledge of genomic signatures underlying host-pathogen relationships, leading to more effective biosecurity and disease management strategies as well as advancements in agricultural pest control (Childers et al. 2021). Further, an increased understanding of genome architecture and functional genomics will shed light into genome size and complexity (Lynch & Conery 2003; Alfsnes et al. 2017) including biological phenomena such as genome streamlining.

## Conclusions

The high-quality genome of *R. argentea*, provides a foundation for functional and evolutionary studies into these tree species, particularly in the context of conservation in the face of myrtle rust. The power of long-read sequencing is further illustrated by the ability to recover a gapless telomere-to-telomere mite genome from contaminating DNA. This study provides points of consideration and recommendations for future genome projects in non-model organisms. Whilst the identity of the contaminating mite remains a mystery, the completeness of its genome should provide an invaluable resource for investigating genome reduction and other characteristic traits of eriophyoid mites.

## Acknowledgements

We thank the horticultural staff at the Botanic Gardens of Sydney with assistance with sampling and Elena Hilario, Cynthia Torkel, and Niccy Aitken for assistance with DNA extraction and MinION sequencing. We are grateful to Jérôme Santolini, Sophie Gerber, and Wannes Dermauw for contacting us about contamination in the draft genome.

## Author contributions

**Stephanie Chen:** Investigation (DNA extraction for long-read sequencing), Methodology, Software, Formal analysis (tree genome assembly and annotation), Writing - Original Draft, Writing - Review & Editing. **Ashley Jones:** Investigation (DNA extraction and MinION sequencing), Writing - Review & Editing. **Patricia Lu-Irving:** Investigation (DNA extraction for 10x linked-read sequencing), Writing - Review & Editing. **Samantha Yap** Investigation (DNA extraction for 10x linked-read sequencing), Writing - Review & Editing. **Marlien van der Merwe:** Investigation and Methodology (DNA extraction for 10x linked-read sequencing), Writing - Review & Editing. **Jason Bragg:** Conceptualization, Writing - Review & Editing, Supervision. **Richard Edwards:** Conceptualization, Methodology, Software, Formal analysis (mite genome analyses and v1 of the *R*. *argentea* genome mentioned in this paper), Writing - Original Draft, Supervision.

## Data accessibility statement

The Rarg_v2 genome (drRhoArge1.2), raw Nanopore long-reads, and Hi-C data were deposited to NCBI under BioProject PRJNA737568. The mite genome (Rarg_mite_v1.1) was also uploaded to this BioProject. The raw 10x linked-reads were deposited under BioProject PRJEB30444 created for an earlier draft genome.

## Benefit sharing statement

All raw sequencing data and assembled genomes have been shared with the broader public via appropriate biological databases.

